# *k*Mermaid: Ultrafast functional classification of microbial reads

**DOI:** 10.1101/2023.08.28.555149

**Authors:** Anastasia Lucas, Daniel E. Schäffer, Jayamanna Wickramasinghe, Noam Auslander

## Abstract

Shotgun metagenomic sequencing can determine both taxonomic and functional content of microbiomes. However, current functional classification methods for metagenomic reads require substantial computational resources and yield ambiguous classifications, limiting downstream quantitative analyses. Existing *k*-mer based methods to classify microbial sequences into species-level groups have immensely improved taxonomic classification, but this concept has not been extended to functional classification. Here we introduce *k*Mermaid, for classifying metagenomic reads into functional clusters of proteins. Using protein *k*-mers, *k*Mermaid allows for highly accurate and ultrafast functional classification, with a fixed memory usage, and can easily be employed on a typical computer.

## INTRODUCTION

Shotgun metagenomics allows for untargeted sequencing of all microbial genomes present in a sample. Unlike 16S sequencing, which is limited to capturing bacterial taxonomy, shotgun metagenomics can be utilized to profile the functional potential of diverse microbiomes. Shotgun metagenomics has been used for identification of microbial biomarkers [1-3], clustering or prediction of disease phenotypes based on microbes [4-6], and for functional analysis [7, 8]. Analyses of shotgun metagenomics can be separated into two major categories: assembly- and read-based, where usage depends on the goals of the experiment. Assembly based analysis first requires read assembly into microbial contigs, whereas read based performs analysis on the unprocessed shotgun reads^8^. While read assembly allows for longer sequences that may be more accurately characterized, de-novo assembly is itself a challenging computational task [9, 10]. Read based analysis overcomes challenges associated with assembly, such as compute time for large volumes of data, coverage of low-abundance organisms, and consideration of substantially mutated or repetitive sequences that are especially difficult to assemble. In addition, read based methods allow for subsequent quantitative analysis, while contigs pose a challenge for read counting [11].

As the field of metagenomics has matured in terms of popularity and technical advancement [12, 13], there has been increasing recognition of the importance of functional analysis [14]. Functional profiling and quantitative comparisons of microbial proteins have immense potential to reveal microbe-microbe and host-microbe interactions, establish new microbial biomarkers and provide predictors based on microbiomes [15, 16]. However, quantitative analysis of microbial functions requires the functional classification of shotgun metagenomic reads, which remains highly challenging. Existing approaches for sequence annotation rely on homology between a query sequence and proteins in a reference databases [16] by aligning the query against a database of microbial proteins. However, metagenomic sequencing experiments typically produce tens of millions of reads [17], rendering the process of read alignment against a typical protein database resource intensive and slow. In addition, most short sequencing reads ambiguously align to multiple microbial proteins from different taxonomies, confounding the ability to count reads and prohibiting downstream quantitative analyses [11, 18, 19].

Several methods have been developed for taxonomic classification of microbial sequences, which have addressed similar challenges in computational efficiency of taxonomic assignment. Most notably, Kraken [20] introduced accurate and highly efficient taxonomic classification by mapping *k*-mers to lowest common ancestors. This approach was later provided with improved resource usage through Kraken2 [21] and improved precision through KrakenUniq [22]. Other high speed taxonomic classification methods are CLARK [23], another *k*-mer based approach, as well as Centrifuge [24] and Kaiju [25], which are based on FM-indexing. Importantly, Kaiju [25] demonstrated that use of protein level sequence comparisons substantially improves taxonomic classification. However, methods offering similar advancement for functional classification are lacking. Whereas mapping reads to proteins using BLASTX [26] is infeasibly slow, DIAMOND [27, 28] allows ultrafast and highly sensitive read-to-protein alignments, making DIAMOND the most widely used tool for functional classification of metagenomic reads [29-31]. However, pairwise alignment which underlies DIAMOND searches does not address ambiguously or multi-mapped sequences and fails to classify sequences with remote homology to proteins in the database. In addition, for typical input metagenomic files, DIAMOND may be resource intensive and require high-performance computing (HPC) or cloud computing systems.

Here, we introduce *k*Mermaid, a new approach for ultrafast and resource efficient functional classification of metagenomic reads (Figure 1). *k*Mermaid unambiguously classifies query sequences into species-agnostic clusters of homologous proteins using a precomputed *k*-mer frequency model. The underlying rationale for *k*Mermaid is that proteins with similar sequences have similar biological function and should be grouped together for downstream analysis. Thus, mapping the sequences to clusters representing homologous groups of proteins irrespective of species addresses issues along both computational and biological axes. First, this approach resolves the problem of alignment ambiguity, defined herein as multi-mapping, when a single read similarly aligns to more than one protein, by collapsing similar protein sequences into more encompassing functional units. Second, by aggregating at the functional level, *k*Mermaid can capture novel biological effects that may be overlooked when performing analysis at the species level. Third, *k*Mermaid can classify tens of millions of sequences in just a few hours providing the computational speed and resource efficiency needed for the large volumes of data generated through metagenomic sequencing experiments while matching the sensitivity of BLASTX. Therefore, *k*Mermaid achieves fast, resource efficient, and sensitive metagenomic read classification into functional units that are expected to substantially improve downstream quantitative analysis.

**Figure 1.**
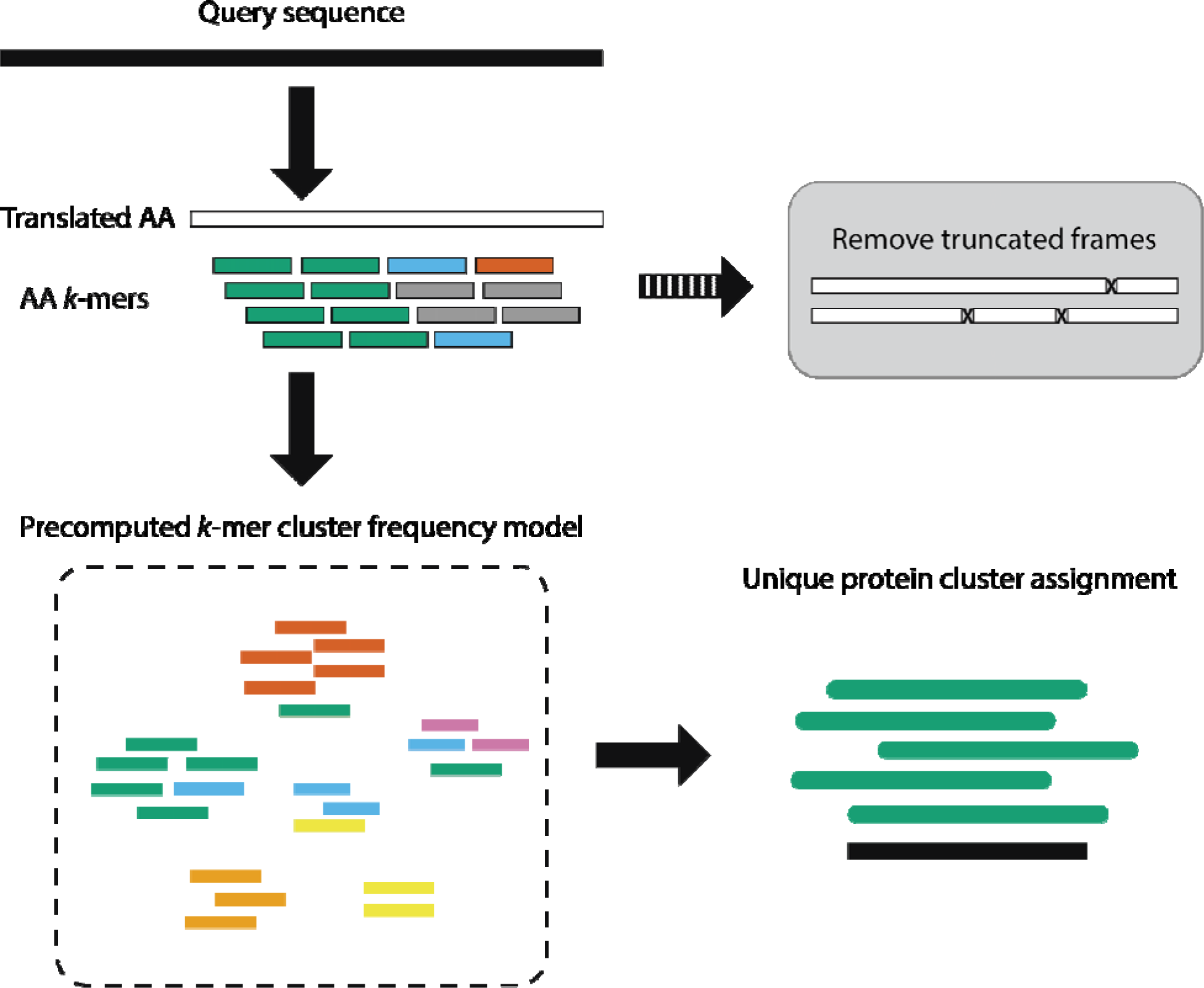
*k*Mermaid metagenomic functional classification. To classify a metagenomic read, the nucleotide sequence is first translated using a six-frame translation, removing the truncated frames from consideration. Each *k*-mer in a non-truncated coding frame is then mapped to the protein clusters that contain that *k*-mer in the database. A composite score is calculated to evaluate a match between the query sequence and each protein cluster, by summing frequencies of *k*-mers in the query sequence in every protein cluster. The query is classified to the protein cluster assigned with the highest composite score (i.e. the cluster in which the *k*-mers of the query are most frequently observed).

## RESULTS AND DISCUSSION

### k-mers clusters of protein functions

The main motivation for *k*Mermaid is that short metagenomic reads are rarely mapped to a single protein and, consequently, a unique read-to-protein mapping requires grouping homologous proteins into functional units. Such functional units become especially critical for downstream quantitative analyses to prevent issues such as double counting multi-mapped reads. We find that most reads are ambiguously mapped by BLASTX with at least 3 protein hits, while only 8% of the reads can be uniquely mapped by alignment to a single protein (Figure 2a). In contrast, by converting the BLASTX hits from proteins to homologous protein clusters (see Methods), we find that more than 95% of the reads can be uniquely mapped to a single cluster or functional unit. In other words, for more than 95% of the reads, all the BLASTX hits of each read belong to a single cluster. This demonstrates that clustering similar proteins together resolves the vast majority of ambiguous BLASTX alignments and in part confirms that protein clusters are able to retain information gained by multiple BLASTX hits.

**Figure 2.**
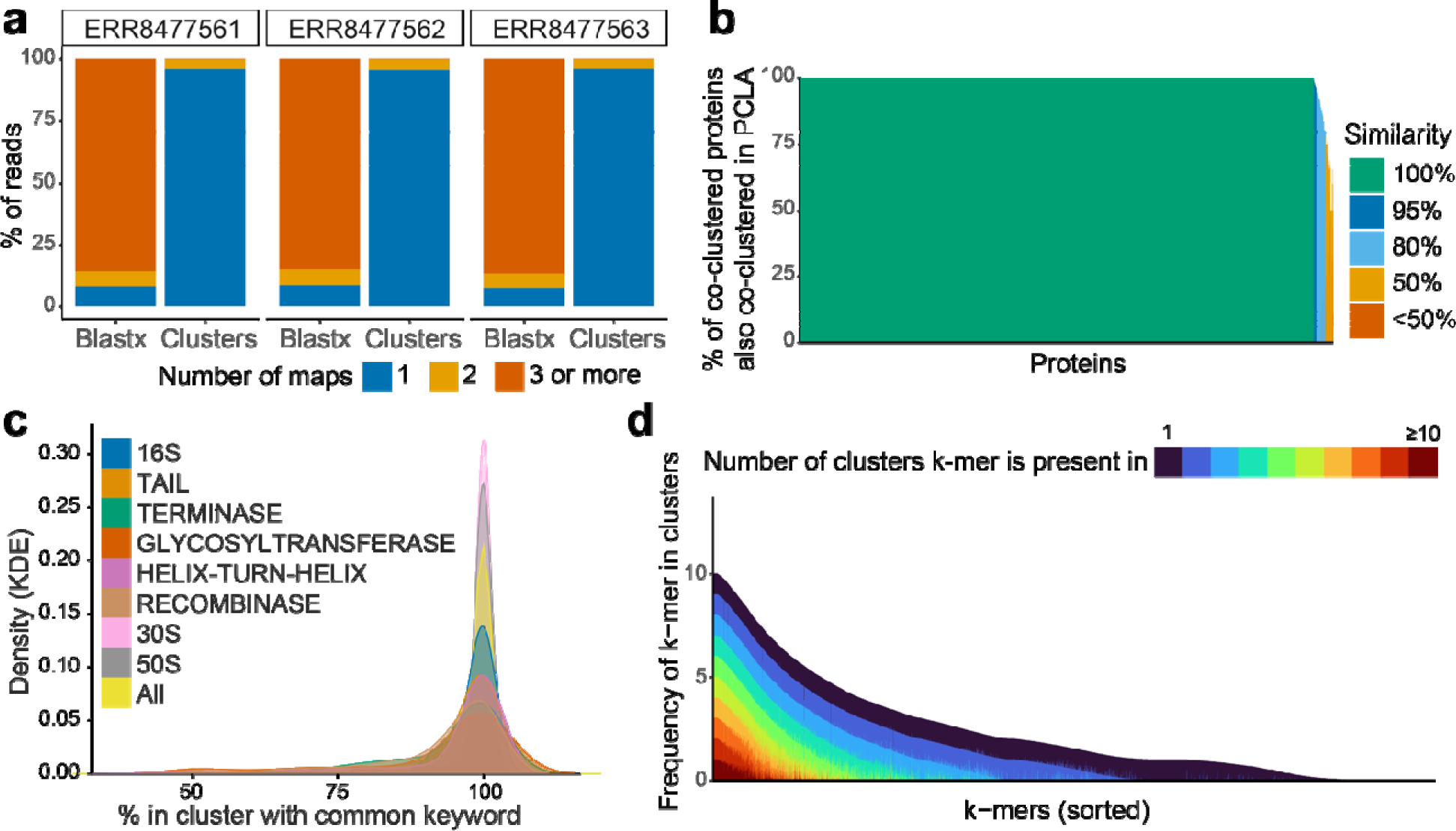
Assessment of the protein clusters underlying *k*Mermaid. **(a)** The reduction in ambiguous maps when looking at read-to-cluster maps compared to read-to-protein level hits (BLASTX). **(b)** The percent of co-clustered proteins also co-clustered in the NCBI PCLA prokaryotic protein clusters for all overlapping proteins were plotted as bars. The x-axis contains individual proteins ordered by cluster, i.e., proteins in the same cluster are plotted next to each other, and the color of the bar corresponds to the percent similarity. **(c)** The distribution (Kernel density estimation, KDE) of keyword percentage, i.e., percent of cluster members with the most common word from all names of proteins in the cluster, across all clusters (yellow) and for clusters with specific common keywords of interest. **(d)** Visual representation of the number of clusters a unique *k*-mer is found in where the y-axis corresponds to the *k*-mer frequency in the cluster. The panel shows a representative, random 50K subset of all *k*-mers.

We ultimately developed *k*Mermaid to efficiently and accurately map microbial short read sequencing to unambiguous homologous clusters of proteins. A user provides a file with sequences to *k*Mermaid to query against our precomputed model of 1,797,426 proteins in 32,308 clusters. For each read in the query file, *k*Mermaid will uniquely assign it into one of the clusters and provide a readable functional annotation based on its cluster representative. Because approximately 25% of our cluster representatives were best given non-descriptive names, *e.g.,* “hypothetical protein,” through RefSeq, we employed HH-suite3 [32] remote homology detection to produce descriptive cluster names. In total, we reannotated 8,617 proteins, of which 6,488 with high confidence (Supplementary Table 1). These include 601 phage proteins, 436 membrane proteins, 230 transcriptional proteins, and 205 lipoproteins (Supplementary Table 1). The composition of *k*Mermaid’s clusters is also highly consistent with preexisting, smaller-scale cluster annotations. We verify that 96% of proteins share 100% similarity with existing NCBI protein clusters, i.e., all proteins in the *k*Mermaid cluster were also co-occurring in the NCBI-derived clusters (Figure 2b). In addition, using keywords of NCBI assigned protein names, we show that the functional annotations of the proteins are highly homogenous within clusters (Figure 2c). Therefore, we comprehensively verify that *k*Mermaid’s cluster naming annotations are biologically accurate and highly reflective of the cluster content.

*k*Mermaid’s internal pipeline assigns sequencing reads to clusters purely based on the composite amino acid (AA) *k*-mer frequencies, where higher confidence scores indicate that *k*-mers within a query sequence are frequently observed in a cluster. This means that for some query sequences, a *k*-mer may uniquely determine cluster assignment, while others may contain multiple *k*-mers that have a higher combined frequency in the assigned cluster compared with any other cluster. With this in mind, we reasoned that a *k*-mer found in many clusters may be less informative and its contribution to the composite score for that cluster is noisy. In the improbable case that all *k*-mers were present in all clusters at a similar frequency, our cluster assignment would be close to random chance. On the other hand, *k*-mers that are only found in one or two clusters would allow deterministic classification and enhance our confidence in use for read assignment. Out of approximately 2.5M AA 5-mers in our model, presence in a single cluster was the most common scenario (22%) and 81% of all AA 5-mers were found in <10 clusters (Figure 2d). We also found that the AA 5-mers that are present in many clusters have relatively similar frequencies across the clusters when examining the top 10 clusters where they are most frequently found. Reassuringly, given the total number of clusters in our model (∼32K), the relative percentage of 5-mers that tend to be deterministic, and the composite score considering multiple *k*-mers in each query sequence, *k*Mermaid clusters fully resolves read alignment ambiguity or multi-mapping in more than 95% of the cases (Figure 2a).

### Performance evaluation

To provide validation for our model, we comprehensively benchmarked the accuracy and sensitivity of *k*Mermaid on both simulated reads and metagenomic sequencing data generated from human samples. We first benchmarked *k*Mermaid on simulated data sets with varying rates of point mutations and single nucleotide insertions or deletions to anticipate how *k*Mermaid would perform on experimental microbial data. Because we wanted to first compare against ground truth, the reads were simulated from the RefSeq data used in the second step of the clustering procedure so that their true cluster labels would be known. Encouragingly, these data showed a high sensitivity (Figure 3a), as *k*Mermaid was able to correctly assign reads to the correct cluster in over 95% of cases across most of the parameters.

**Figure 3.**
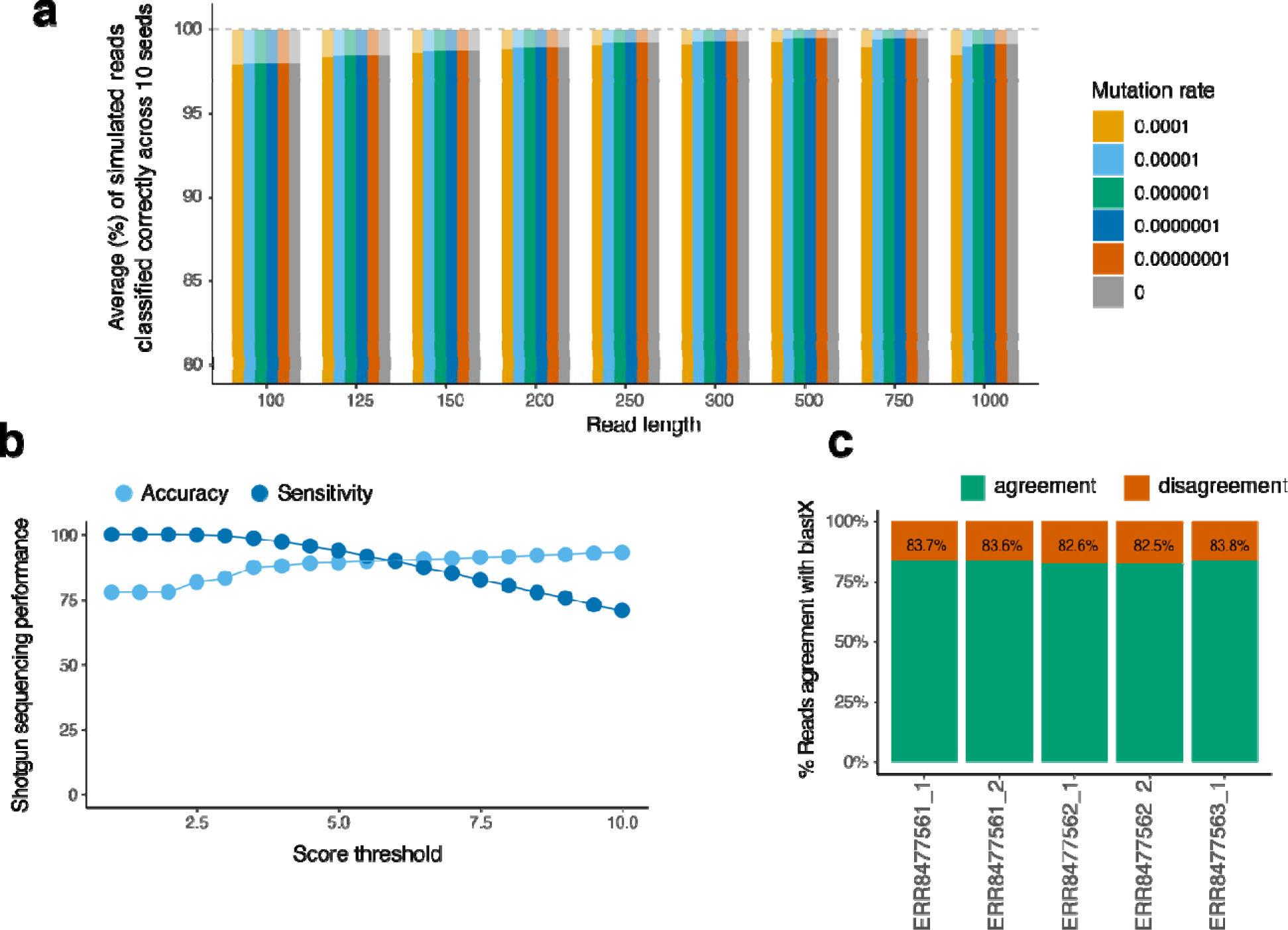
Benchmarking *k*Mermaid’s performance on simulated and human fecal data. **(a)** The percentage of reads of classified correctly by *k*Mermaid across 10 simulated datasets per each combination of read length and mutation rate (opaque bars) out of the percentage of reads that were able to be classified (bars with transparency) is between 92-99% and correlates with mutation rate. **(b)** The percent of all input reads able to be classified by *k*Mermaid compared to BLASTX for sequencing from 5 fecal samples. *k*Mermaid’s optimal scoring threshold was determined by maximizing the percentage of the assignments that agree with BLASTX hits (Sensitivity, dark blue bars) while retaining a high ratio of assignments that agree with BLASTX to assignments that disagree with BLASTX (Accuracy, light blue bars). **(c)** *k*Mermaid consistently achieves around 83% agreement (green) with BLASTX.

Even though *k*Mermaid performed well on simulated reads, by simulations alone it is impossible to account for the additional challenges and noise associated with real, experimental data. Therefore, we performed additional benchmarks using real sequencing data from publicly available human fecal samples, comparing *k*Mermaid assignment against BLASTX. Assuming BLASTX hits to be the ground truth, *k*Mermaid was able to maintain a balance between retaining a high percentage of the assignments that agree with BLASTX hits (referred to as sensitivity) as well as a high ratio of assignments that agree with BLASTX to assignments that disagree with BLASTX (referred to as accuracy) (Figure 3b) at a *k*Mermaid confidence score ≥ 3. On average, *k*Mermaid was able to map 25% of the input reads to a protein cluster compared to 4-5% classified by BLASTX. *k*Mermaid’s BLASTX coverage (the percentage of BLASTX classified reads classified with *k*Mermaid) was consistently around 81% for the samples analyzed. These high confidence hits also showed consistent agreement with BLASTX, with around 83% of the reads having overlap between the assignment of the two approaches (Figure 3c). Together with the result that the majority of a read’s BLASTX hits belong to a single cluster (Figure 2a), these findings provide strong evidence for *k*Mermaid’s ability to accurately and sensitively classify short reads from metagenomic sequencing data.

### Computation efficiency

Shotgun metagenomics experiments commonly yield tens of millions of sequences per sample and each must be queried against a large reference database for classification purposes. BLASTX remains the gold standard for this task, but its computational time is infeasible for typical large files, necessitating the development of alternatives that can process these reads in a reasonable timeframe. We benchmarked *k*Mermaid’s runtime against BLASTX and DIAMOND [27, 28], which, to our knowledge, is the fastest approach for functional annotation of metagenomic reads and was in part developed to address the runtime limitations of BLASTX. *k*Mermaid matches the ultrafast running time of DIAMOND and both methods ran from 10 to 25 times faster than BLASTX on files with up to 50M sequences. Both *k*Mermaid and DIAMOND were able to run 500K sequences in a minimum of 3.5 minutes and 4 minutes, respectively. The same input when given to BLASTX took over 6 days to complete. Further, classifying 50M sequences took both *k*Mermaid and DIAMOND between 5 and 6 hours on average, which highlights their usability for real shotgun metagenomic sequencing files. Additionally, since *k*Mermaid classifies reads independently of other reads in the same input file, it easily lends itself to parallelization by means of splitting input files into smaller chunks, allowing for further speed improvement when resources are available.

Along with speed, RAM usage is another potentially limiting factor when input files are large. We therefore compared *k*Mermaid to DIAMOND, which has excellent running times, but achieves this performance in speed at the expense of high memory and multiple CPU utilization. With these limitations in mind, we have developed *k*Mermaid to be highly memory efficient. *k*Mermaid perform read assignments in a way that only requires the precomputed k-mer frequency model, and not the input data, to be loaded into memory. In this way, *k*Mermaid requires a fixed amount (2GB) of memory per run regardless of file size, whereas DIAMOND which requires memory to scale with increasing input file size with a cap of 16G RAM (Figure 4c).

**Figure 4.**
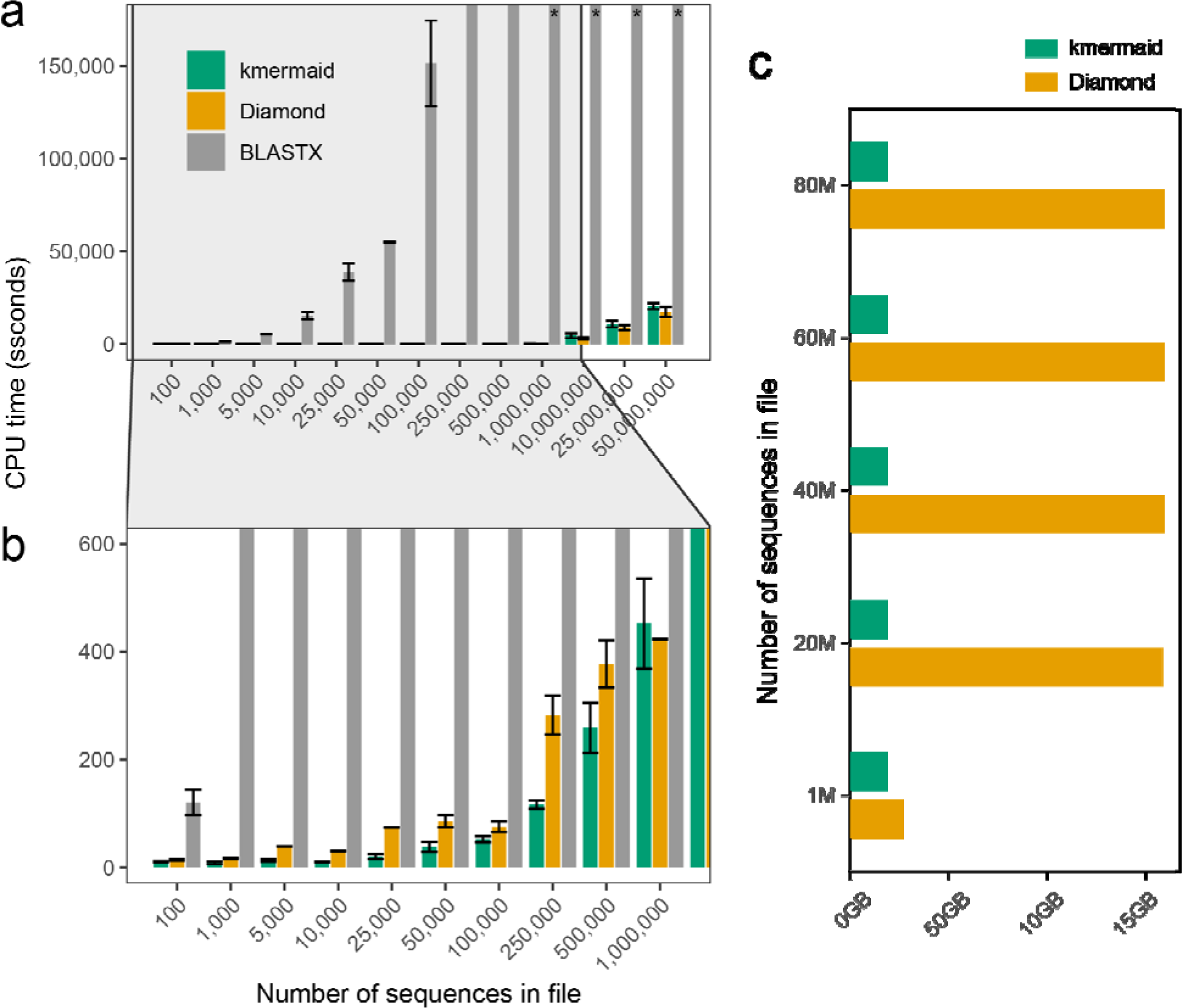
Runtime and memory usage comparisons to BLASTX and DIAMOND. **(a)** *k*Mermaid (green) provides up to 25-fold decrease in runtimes compared to BLASTX (gray, files that exceeded a 7-day run time denoted with an asterisk) and has comparable runtimes to DIAMOND (orange). The y-axis has been truncated due to excessive running times of BLASTX. **(b)** A zoomed-in panel showing that 500K sequences can run in <10 minutes using DIAMOND and *k*Mermaid. **(c)** *k*Mermaid (green) requires a fixed, low memory allotment to run while DIAMOND’s (orange) RAM utilization scales with file size. BLASTX was excluded from this comparison due to the infeasible running times for files of this size.

### Cluster specific results

The primary goal of *k*Mermaid is to achieve high performance for classification at the read level. To comprehensively assess *k*Mermaid’s performance, we also evaluated the cluster specific performance of *k*Mermaid. We compared the *k*Mermaid read assignments of the real metagenomic sequencing input samples used for benchmarking to the assignment by BLASTX, within each cluster. Interestingly, we find that clusters related to restriction, toxins, transposons, and those in the GCN5-related N-acetyltransferases family (GNAT) had high agreement with BLASTX, whereas clusters related to ABC transporters tended to have relatively low agreement with BLASTX, and therefore likely lower accuracy (Figure 5a). We also confirmed that the proportion of reads concordant with BLASTX was correlated with the mean *k*Mermaid assigned confidence score for all reads mapping to the cluster, a trend that was not confounded by the number of reads mapped to the cluster (Figure 5b). Importantly, by investigating reads that were assigned with a high *k*Mermaid confidence score but were not classified by BLASTX, we identified reads with remote homology to proteins within *k*Mermaid clusters. We verified a correct functional classification of reads assigned to six such *k*Mermaid clusters using both PSI-BLAST [33] and HHblits3 from HH-suite3^24^ and validated these *k*Mermaid functional annotations (Figure 5c, Supplementary File 1).

**Figure 5.**
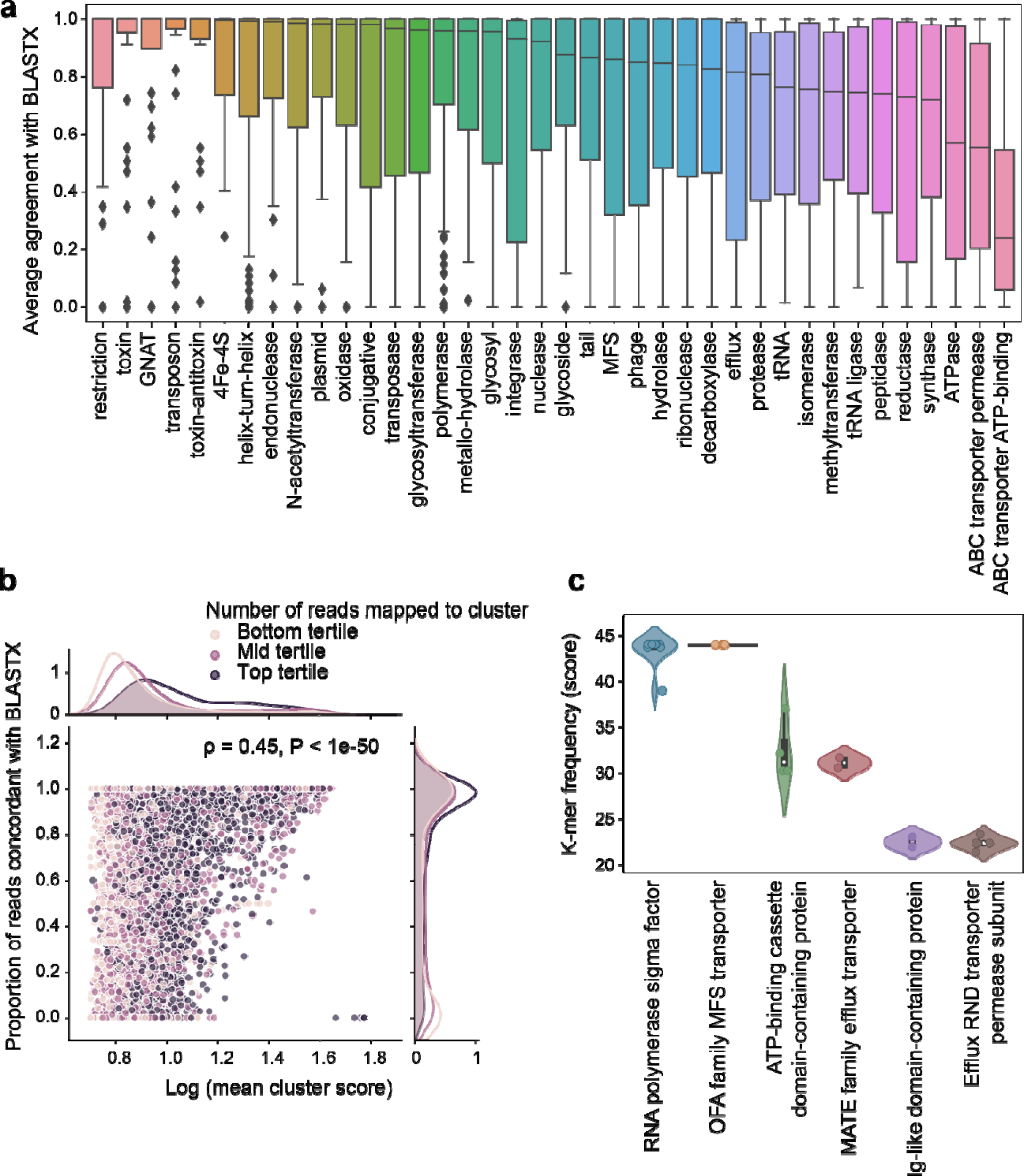
Function specific performance and evidence of remote homology detection. **(a)** Boxplot showing *k*Mermaid performance (agreement with BLASTX) for clusters with specific functional annotations. **(b)** Correlation between the proportion of reads concordant with BLASTX and the mean confidence scores (log-transformed) for all proteins in the cluster. Distributions of these metrics broken down by number of reads mapped to the cluster where clusters in the bottom tertile have the lowest number of mapped reads and clusters in the top tertile contain the highest. **(c)** violin plots showing the *k*-mer frequency scores for six clusters of reads unclassified with BLASTX that were correctly functionally classified by *k*Mermaid.

## CONCLUSIONS

Metagenomic sequencing allows for the functional profiling of diverse microbes facilitating numerous biomedical applications, such as biomarker discovery and disease prediction [34-36]. To date, *k*-mer and binning based methods have been immensely useful in allowing efficient and sensitive classification of short read metagenomic sequencing into taxonomic units [20-23, 37-40]. However, to the best of our knowledge, *k*-mer or binning based methods for functional classification have not been developed. As a result, there remain critical limitations in our ability to functionally classify short microbial reads, which are typical to Illumina next generation sequencing platforms. The main limitations are ambiguous read alignments and computational costs that are prohibitive for the large volumes of data typically generated by sequencing experiments. As such, there is a need for methods that can capture and retain the underlying biology of microbial function in a computationally feasible and accessible manner.

To address these challenges, we have developed *k*Mermaid, a novel ultrafast method for unambiguous and sensitive classification of short reads into functional units consisting of protein clusters. We show that by considering a cluster of homologous proteins as a single functional unit, *k*Mermaid resolves the majority of ambiguous BLASTX proteins alignments while retaining the granular functional information. *k*Mermaid uses a novel precomputed *k*-mer frequency model based on high-confidence protein clusters encompassing almost 2M microbial proteins from RefSeq. Using both simulated short reads and sequencing data from real human fecal samples, we demonstrate that *k*Mermaid classifies reads with high accuracy and sensitivity compared to BLASTX but runs up to 2,500% faster on typical files with of tens of millions of sequences. Additionally, we were able to verify *k*Mermaid’s ability to correctly classify reads that were unclassified by BLASTX for sequences with remote homology to proteins within *k*Mermaid clusters. This striking result is likely allowed through *k*Mermaid’s composite scoring which uses information on a set of proteins in a cluster to classify each read in contrast to BLASTX, which is based on pairwise comparisons.

This study has several potential limitations that should be acknowledged. First, while *k*Mermaid assigns a read to a protein cluster based on a global maximum *k*-mer frequency score, in the case of ties it will randomly assign the sequence to a cluster based on the first time it reaches the maximum. Despite this, we have shown that ties should not be a widespread issue since the vast majority of multi-mapped reads are mapped to a single cluster (Figure 2a), and many AA k-mers are unique (Figure 2d). Second, despite our efforts to compare the performance of *k*Mermaid on experimental data to a ground truth, extensive and large-scale manual curation would be required to compute the true accuracy of *k*Mermaid beyond what we can determine by comparison with BLASTX. Third, it is possible that additional clusters aside from the ones provided through *k*Mermaid database exist which are not homologous or functionally similar. As such, it is recommended that researchers use our method as a first pass means of dealing with the computational burden of BLASTX functional classification and perform alignment-based verification of functions of interest.

In summary, we present *k*Mermaid, a novel, sensitive, runtime and memory efficient approach for the task of assigning function to short microbial sequences. Future studies can utilize *k*Mermaid for the discovery of microbial functional biomarkers and as a precursor to downstream quantitative functional analyses.

## METHODS

### Establishing clusters of functionally similar microbial proteins

*k*Mermaid is foremost designed to address ambiguity in read alignment, where most shotgun metagenomic reads align against multiple microbial proteins with similar alignment scores and are therefore classified as multiple functionally-related proteins by the alignment (Figure 2a). To circumvent this issue, the first goal of *k*Mermaid is to group functionally related proteins into functional units prior to assignment such that a read can be uniquely classified into a single functional unit. We therefore constructed a set of comprehensive and high confidence clusters of microbial proteins by employing a two-step clustering procedure using CD-HIT[41]. CD-HIT uses an incremental greedy algorithm to identify representative sequences and cluster remaining sequences by sequence similarity using short word filtering. We first clustered 43,176 proteins from the NCBI RefSeq representative microbial protein database using CD-HIT with a similarity threshold of 65% and a word size of 5. This first clustering step resulted in 32,308 clusters, including clusters of single proteins that have <65% sequence similarity with any other cluster representative. In the second step, to further expand and diversify these clusters, we applied CD-HIT-2D[42] with a 70% sequence similarity threshold to cluster all RefSeq microbial proteins (N=1,797,426, as of May 2023) against the previously selected cluster representatives. A set of expanded clusters was created from this process such that the final dataset clusters 1,793,361 RefSeq proteins into 32,308 functional groups.

### Evaluating the clusters of functionally similar microbial proteins

As the protein clusters lie at the base of *k*mermaid’s approach, we validated their correctness using two analyses aimed to verify that the clusters produced contain homologous proteins with shared biological function.

1. *Compatibility with NCBI protein clusters.* To demonstrate *k*Mermaid’s ability to construct biologically relevant clusters, we compared the results of the two-step CD-HIT clustering to the datasets from the NCBI Protein Clusters[43], which groups together proteins by sequence similarity. A subset of 102,380 of the proteins contained in our expanded cluster model were also clustered through the prokaryotic PCLA protein clusters dataset within this database. Proteins that were in this overlapping subset and were also in a non-singleton *k*Mermaid cluster were used for comparison (N = 102,367, mapped to 9,984 *k*Mermaid clusters). For each of these proteins, we evaluated its tendency to be co-clustered with the same proteins in both PCLA and *k*Mermaid clusters by computing the percent of co-clustered proteins by *k*Mermaid clustering that are also co-clustered in PCLA. The number of *k*Mermaid clusters was chosen as the denominator for this evaluation metric to verify the correctness of *k*Mermaid clusters rather than to assess its ability to maximize clusters, which is not an objective of this approach.

2. *Within-cluster keyword similarity.* High-throughput text analysis was performed on the protein name annotations to further investigate the similarity and homogeneity of the clusters. Trends and frequencies of word presence in clusters were used to evaluate cluster functional homogeny. After removing ubiquitous and generic words (e.g., “bacteria” or “protein”), we computed the frequency of the most common keyword found in each cluster, i.e., the fraction of proteins in the cluster containing the most common keyword in that cluster. The majority of clusters demonstrated a common keyword frequency of 1, indicating that our clusters are highly homogenous in key functions.

### Assigning functions to clusters of hypothetical or uncharacterized microbial proteins

*k*Mermaid annotates protein clusters with representative names derived from the functional annotation of the protein mapped to each cluster’s representative sequence, determined in the initial CD-HIT phase on the representative protein database. Even though these are in the representative microbial protein database, 8,617 (approximately 25%) of the cluster representative are best annotated by NCBI or RefSeq as hypothetical proteins, i.e. proteins of unknown or unverified function. To annotate these protein clusters and assign them with functional description, we used HHblits from HH-suite3 [32] for remote homology detection against two databases from The Protein Databank and UniProtKB (PD70 [44] and Uniclust30 [45] v2023, respectively) and selected the match with lowest e-value across the databases that did not map to hypothetical, unknown, or uncharacterized proteins. We were able to confidently assign function to 6,488 of the 8,617 hypothetical clusters (with e-value < 0.01) and the rest are assigned with lower confidence.

### k-mer frequency-based protein cluster assignment model

The ultimate goal of *k*Mermaid is to functionally classify short reads using the previously defined clusters of functionally similar microbial proteins. To this end, we built a *k*-mer frequency model by obtaining all amino acid (AA) *k*-mers of all protein sequences in each cluster and computing the cluster-level frequency of each AA *k*-mer. This frequency is defined by the count of the *k*-mer in the cluster divided by the total number of proteins in the cluster. This precomputed model, consisting of 2,574,615 unique *5*-mers, is then used to determine the cluster to which a query sequence is assigned.

### Functional classification of reads using the pre-computed k-mer model

To assign a read into protein cluster, six-frame translation is applied followed by removal of truncated frames. Next, all AA *k*-mers are extracted from non-truncated translated frames. A composite score that represents the strength of a match between a read and each cluster is then calculated by the summation of the precomputed model *k*-mer frequencies for the set of *k*-mers in the query sequence. The sequence is then assigned to the cluster with the global maximum composite score. This approach effectively assigns higher confidence scores when *k*-mers within a query sequence are more frequently observed in a cluster. *k*Mermaid uses a *k* of 5 AA chosen via hyperparameter search for both the base model construction and the assignment procedure and provides assignments with confidence score > 3.

### Performance assessment using simulated reads

To demonstrate that *k*Mermaid correctly assigns proteins to clusters, we benchmarked *k*Mermaid using data simulated from 1,383 microbial coding frames downloaded from RefSeq for which the true cluster identity is known, i.e., proteins that already exist in *k*mermaid’s model and thus have a ground-truth cluster label. From these, simulated data were generated with varying mutation rates, mutation types, and read lengths (Table S1). Mutation rates were chosen to be representative of bacterial mutation rates [46] and error rates in next-generation sequencing data [47]. The number of mutations per sequence was determined probabilistically using a binomial distribution and the location of the mutation in the sequence was determined by random sampling. Point mutations were defined as a random assignment of any nucleotide that did not match the original position; indels were assigned similarly and determined to be either single nucleotide insertions or deletions at random. Query sequences were then created by segmenting the mutated sequence to the specified read length and assigned to clusters by *k*Mermaid as described above. Results from the simulations were averaged across 10 replicate datasets generated with a different random seed for each combination of parameters.

### Performance assessment using real metagenomic data

The goal of *k*Mermaid is to efficiently map reads to proteins, while maintaining the accuracy and sensitivity of BLASTX. We benchmarked *k*Mermaid’s functional read classification against BLASTX using paired end reads from ulcerative colitis patients enrolled in the LOTUS fecal matter transplant clinical trial [48] available on the NCBI’s Sequence Read Archive (SRA). We used a small subset (n=3) of the available samples due to the infeasibly long running times incurred by BLASTX. For each of the fastq files, we ran *k*Mermaid and BLASTX using an e-value cutoff of 0.01 and similarity >66%. We compared the overall coverage, defined as the percentage of reads that were able to be classified by both BLASTX and *k*Mermaid, as well as the percentage of BLASTX hits that *k*Mermaid was able to classify. We further investigated the correctness of *k*Mermaid’s assignments using BLASTX results as a gold standard by looking at the reads which received an assignment by both methods. Because BLASTX can assign a read to multiple proteins with similar scores whereas *k*Mermaid produces one cluster assignment per read, the case when at least one of a read’s BLASTX hits was a member of the *k*Mermaid assigned protein cluster was considered to be a correct assignment by *k*Mermaid.

### Computational efficiency comparison of kMermaid with BLASTX and DIAMOND

We compared the speed of *k*Mermaid to BLASTX and DIAMOND[27, 28], a highly efficient and sensitive protein aligner. Although the goals of *k*Mermaid and DIAMOND are different, DIAMOND, which is up to 20,000 times faster than BLASTX, is the fastest protein aligner that can maintain the sensitivity of BLASTX results and is included in our benchmarking as a standard for efficient resource consumption. To compare the runtime of each method, we created random subsets of a single fasta file containing paired end sequencing data also from the LOTUS clinical trial. For running time comparisons, we tested input files with a varying number of sequences ranging from 100 to 50M with 10 replicates each to account for machine or algorithmic variability. BLASTX and DIAMOND sequence queries were performed against the same database that was used to create *k*Mermaid model described above. All comparisons were run on a Linux kernel using 1 task and 1 CPU per task and CPU time in seconds was extracted after jobs completed using the Slurm seff command. Because it can take BLASTX between weeks and months to run millions of sequences, we set an upper time limit of 7 days and denote jobs that were unable to be completed in that time frame.

### Memory usage comparison between kMermaid and DIAMOND

Individual metagenomic sequencing experiments can yield large volumes of data and file sizes are commonly on the order of tens of gigabytes. As such, efficient memory usage is another important factor to consider when choosing analysis tools. We compared the memory (RAM) usage between *k*Mermaid and DIAMOND for input files containing 20M, 40M, 60M, and 80M reads, which corresponds to the total number of reads commonly generated from a standard paired-end sequencing experiment [46] (10-40M each, combined paired end), as well as a small input of 1M reads. BLASTX was excluded from this comparison due to its infeasibly high running time, although it is noted that BLASTX memory usage is minimal.

### Cluster-specific results and remote homology detection

To evaluate *k*Mermaid’s performance for specific biological functions, we have calculated the accuracy across clusters with similar functional annotations. We used the LOTUS clinical trial data to evaluate function specific performance in a real metagenomic sequencing cohort. To this end, for every cluster we calculated the ratio of reads correctly assigned to that cluster (i.e. assigned to that cluster by BLASTX and *k*Mermaid), out of all the reads assigned to that cluster by *k*Mermaid. Then, we evaluated the distribution of cluster performances for distinct functions, i.e., clusters named with common keywords (Supplementary Table 2).

To evaluate *k*Mermaid’s ability to identify sequences with remote homology to proteins in the database, we examine reads that were not classified by BLASTX from the LOTUS clinical trial data and were assigned with high *k*Mermaid confidence scores (>20). We explored reads that were confidently assigned to six clusters which failed to be classified with BLASTX, and carefully verified that these reads have remote homology to their *k*Mermaid assigned clusters using PSI-BLAST [33] and HHblits3 from HH-suite3^24^ (Supplementary File 1).

### Availability and usage

*k*Mermaid is freely available as an open-source command line program written in Python that requires a user-provided input file containing query sequences in either fastq or fasta format. The precomputed *k*-mer frequency model and files containing the protein cluster members and cluster representatives are both accessible and used as internal default parameters in *k*Mermaid. Complete download and installation instructions are found at github.com/AuslanderLab/kmermaid.

### Availability of data and materials

All data is used in this work is publicly available. The LOTUS trial paired end sequencing metagenomic reads from ulcerative colitis patients fastq files used for benchmarking are available through SRA: PRJEB50699.

### Author Contributions

N.A. initiated the project and supervised work. A.L., D.E.S., J.W., and N.A wrote and tested the software. A.L. and N.A. designed and performed analyses and benchmarking comparisons.

